# A drug candidate for treating adverse reactions caused by pathogenic antibodies inducible by COVID-19 virus and vaccines

**DOI:** 10.1101/2021.07.13.452194

**Authors:** Huiru Wang, Xiancong Wu, Yuekai Zhang, Qiuchi Chen, Lin Dai, Yuxing Chen, Xiaoling Liu

## Abstract

In a recent study, we reported that certain anti-spike antibodies of COVID-19 and SARS-CoV viruses can have a pathogenic effect through binding to sick lung epithelium cells and misleading immune responses to attack self-cells. We termed this new pathogenic mechanism “Antibody Dependent Auto-Attack” (ADAA). This study explores a drug candidate for prevention and treatment of such ADAA-based diseases. The drug candidate is a formulation comprising N-acetylneuraminic acid methyl ester (NANA-Me), an analog of N-acetylneuraminic acid. NANA-Me acts through a unique mechanism of action (MOA) which is repairment of the missing sialic acid on sick lung epithelium cells. This MOA can block the antibodies’ binding to sick cells, which are vulnerable to pathogenic antibodies. Our *in vivo* data showed that the formulation significantly reduced the sickness and deaths caused by pathogenic anti-spike antibodies. Therefore, the formulation has the potential to prevent and treat the serious conditions caused by pathogenic antibodies during a COVID-19 infection. In addition, the formulation has potential to prevent and treat the adverse reactions of COVID-19 vaccines because the vaccines can induce similar antibodies, including pathogenic antibodies. The formulation will be helpful in increasing the safety of the vaccines without reducing the vaccine’s efficacy. Compared to existing antiviral drugs, the formulation has a unique MOA of targeting receptors, broad spectrum of indications, excellent safety profile, resistance to mutations, and can be easily produced.

## Introduction

Many patients with the coronavirus disease 2019 (COVID-19) lost their lives due to unclear understanding of the disease’s pathogenic mechanism and the lack of effective medicines for the disease^1^,^2^. It has also been reported that COVID-19 symptoms can last weeks or months for some recovered patients termed “long haulers”^3^. There are yet no effective treatments for the symptoms of COVID-19 long haulers. Vaccines are the most effective intervention to control the COVID-19 pandemic^4^. However, vaccines are not perfect as they may cause serious adverse reactions and even death^5^. Treatment for the adverse reactions of vaccines will be helpful in eliminating the concerns about the COVID-19 vaccines’ safety and will help promote the increased vaccination globally.

Our recent study reported a newly discovered pathogenic mechanism of COVID-19 infection, one that is induced by certain anti-spike antibodies of the COVID-19 virus^6^. These antibodies are pathogenic in of themselves, and can target and bind to host vulnerable cells or tissues, initiate a self-attack immune response, and lead to serious clinical conditions including acute respiratory distress syndrome (ARDS), cytokine storms, and death^6^. We have termed this novel mechanism “Antibody Dependent Auto-Attack” (ADAA) in a recent article. Since anti-viral antibodies continue to exist for a period of time in recovered COVID-19 patients, certain pathogenic antibodies among them can induce the symptoms of long haulers^6^. Given the similarities of antibodies inducible by COVID-19 vaccines, the pathogenic anti-spike antibodies, despite only being minority of the antibodies induced, provided a possible cause for the adverse reactions of COVID-19 vaccines. Therefore, a treatment capable of blocking the binding of pathogenic antibodies to host cells will be effective for the ADAA-based diseases, including adverse reactions of vaccines.

Sialic acids are rich on the outer surface of cell membranes and mainly act as biological masks^8^. Cells or tissues with sialic acid are recognized as “self”. After the loss of sialic acids the cellular structures become “non-self” ^8^, which can activate immune responses. In a recent report, we have shown that damaged lung epithelium cells with missing sialic acid and human inflammatory tissues are particularly vulnerable to pathogenic anti-spike antibodies^6^. During the COVID-19 infection, the sialic acid on lung epithelium cells can be removed by the receptor-destroying enzyme (RDE) of SARS-CoV-2 virus^9^, that makes the damaged cells vulnerable to pathogenic antibodies. Thus, repairment of the missing sialic acid can convert the sick cells (non-self) back to normal (self) and block the binding of pathogenic antibodies to those vulnerable cells. This newly discovered mechanism of action (MOA) provides a possible new direction for developing anti-viral drugs. Therefore, we designed a formulation consisting of an analog of sialic acid for repairing the missed sialic acid on the vulnerable cells. In this study, we tested the formulation’s efficacy in preventing and treating adverse reactions caused by pathogenic anti-spike antibodies of COVID-19 and SARS-CoV viruses.

## Results

### The composition of a formulation

A successuful drug for repairment of missing sialic acid on the cell surface should have three attributes: 1) easily enter cells; 2) participate in the sugar chain synthesis process without affecting the glycosidic linkages; and 3) be stable *in vivo*. Sialic acid has a negative charge on its surface and cannot easily enter cells and participate in the sugar chain synthesis process, and is metabolized rapidly *in vivo* ^8^. Therefore it is difficult for sialic acid alone to be used as a therapeutic medicine.

N-acetylneuraminic acid methyl ester (NANA-Me) is an analog of N-acetylneuraminic acid (NANA) with a structure close to the structure of NANA. It is easier for NANA-Me to enter cells compared with NANA. However, NANA-Me is unstable at about pH 7.0 (unpublished data). Our study found that the NANA-Me was more stable under the pH conditions of 4.0-5.5 (unpublished data). We further found that the stability of NANA-Me was better when it was in a composition comprising NANA under certain ratio range (NANA-Me:NANA, 1-5:1) of the two compounds (unpublished data, patent pending). Therefore, we designed a formulation consisting of NANA-Me and NANA with a ratio of NANA-Me:NANA at 2:1 (pH 4.43). The formulation was named as *BH-103* (patent pending).

### Repairment of cell surface sialic acid by NANA-Me

The sialic acid repairment function of NANA-Me was tested by a cellular assay. Lung epithelium cell line A549 was cultured with NANA-Me or NANA with various concentrations for overnight, and the sialic acid levels on the cells’ surfaces were determined next day with fluorescent labeled-wheat germ agglutinin (WGA) (Vector) and flow cytometry as described in methods. The sialic acid levels of the A549 cells treated with NANA-Me increased in a way of dose dependent between the range of 1μg/ml - 50 μg/ml, while the sialic acid levels of the A549 cells treated with NANA were not changed significantly (FIG 1A and 1B). The data suggests that NANA-Me is helpful in the synthesis and expression of the sialic acid on lung epithelium cells.

**Figure 1.**
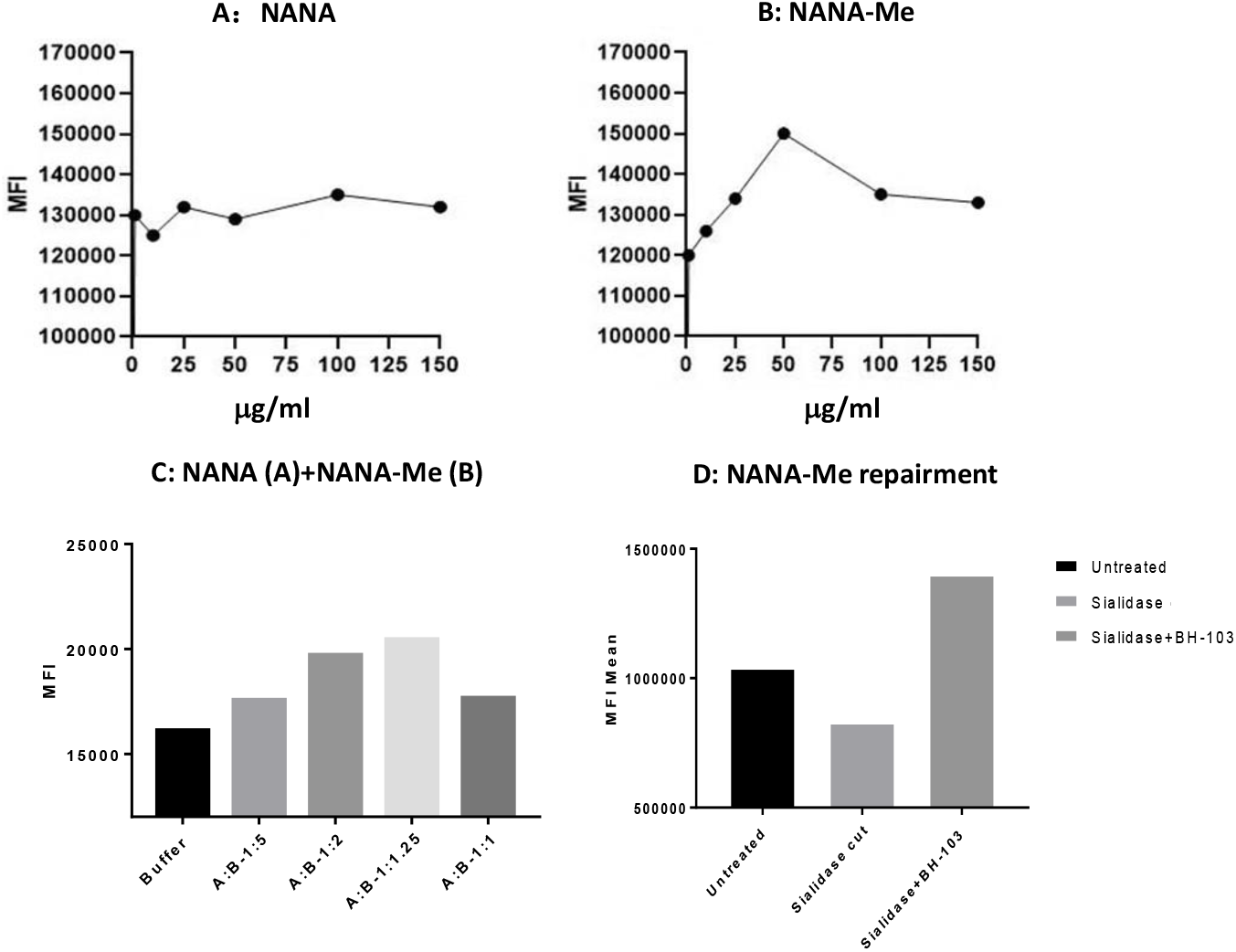
The sialic acid levels on the surface of human lung epithelium A549 cells determined by a flow cytometry analysis. A549 cells were treated with A: N-acetylneuraminic acid (NANA); B: N-acetylneu-raminic acid methyl ester (NANA-Me); and C: NANA+ NANA-Me. D: BH-103 treated A549 cells with missing sialic acid on the cell surface (sialidase treated), compared to control cells of healthy (intact) or damaged A549 cells (sialidase treated) alone.

In another test, the A549 cells were cultured with 50 μg/ml of NANA-Me in combination with NANA at various ratios (NANA-Me:NANA) for overnight. The sialic acid levels on the cells were determined next day with the same method as described above. The results showed that the sialic acid levels on the A549 cells treated with the compositions comprising NANA-Me and NANA at the ratios of 1.25:1 or 2:1 (NANA-Me:NANA) were higher compared with the buffer (vehicle) treated control cells (FIG 1C). Thus, a formulation consisting of NANA-Me and NANA at a ratio of 2:1 (pH 4.43 when dissolved), was prepared and named as BH-103.

In order to induce damaged cells, the A549 cells were treated with neuraminidase or sialidase (Roche, Shanghai) according to manufacturer’s instructions ^6^. The sialidase treated A549 cells were cultured without or with 50 μg/ml of BH-103 (NANA-Me) at 37°C for overnight, and the sialic acid level on A549 cells were determined the next day. As shown in FIG 3D, the sialic acid levels of the sialidase treated A549 cells were lower than that of the untreated cells, indicating loss of sialic acid on A549 cells after being digested with sialidase. The sialic acid levels of the A549 cells treated with sialidase and BH-103 was higher than that of control cells treated with sialidase but without BH-103. Taken together, the data of the *in vitro* analysis indicated that N-acetylneuraminic acid methyl ester has the potential to enhance the expression of sialic acid and to repair the missed sialic acid on the A549 cell surface. The best enhancing or repairing effect of NANA-Me could be achieved in combination with NANA (pH 4.43). The repairment of sialic acid on cell surfaces is helpful for recovery of damaged (e.g. infected or inflamed) cells, blocking the self-attacking response of the immune system and reducing the severity of diseases and preventing deaths.

**Figure 2.**
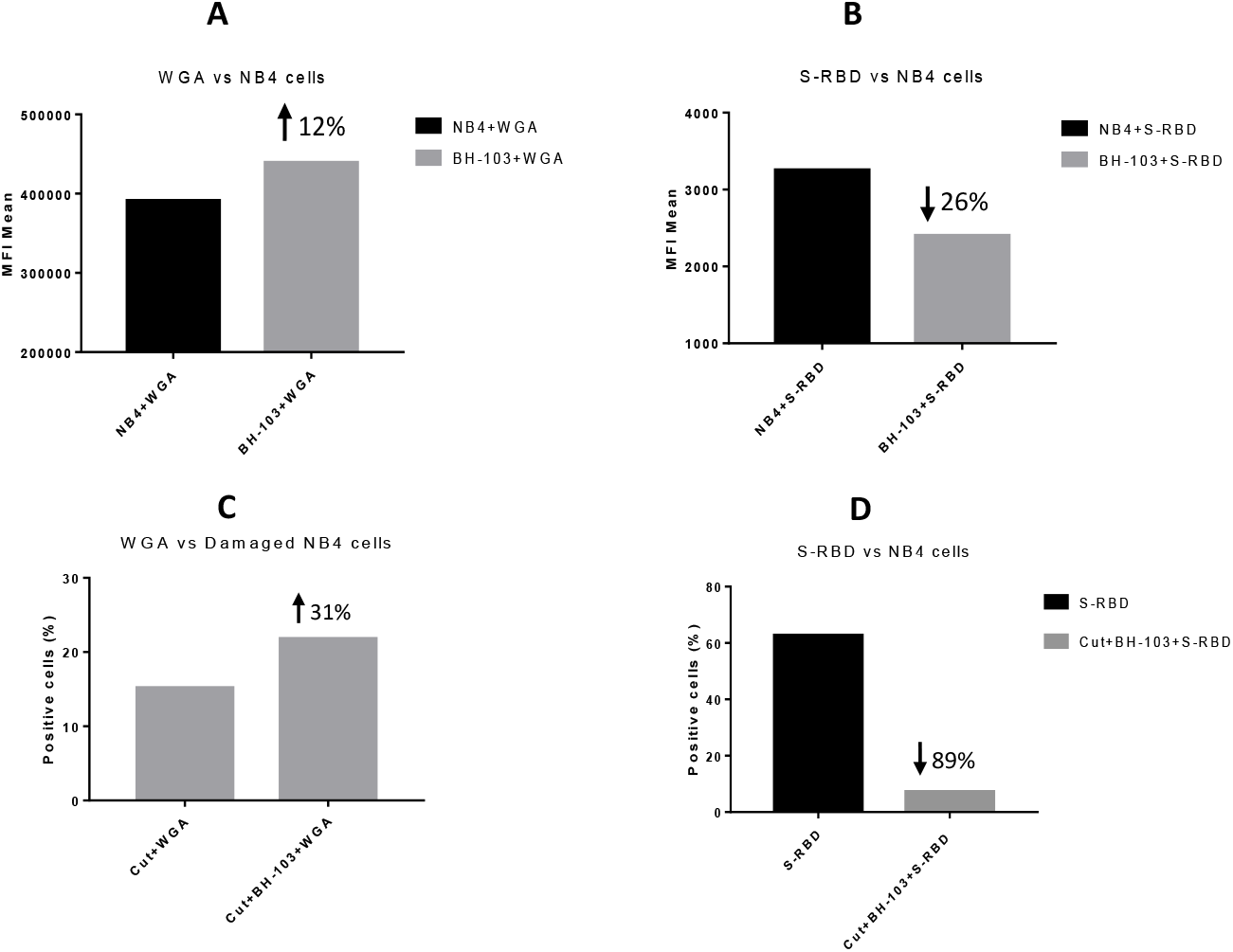
The levels of sialic acid or S-RBD of SARS-CoV-2 virus on the surface of acute promyelocytic leukemia NB4 cells, determined by a flow cytometry analysis. A: NB4 cells were treated with or without BH-103; B: Binding of S-RBD to NB4 cells with or without BH-103 treatment; C: Damaged NB4 cells (sialidase treated) were treated with or without BH-103; and D: Binding of S-RBD to damaged NB4 cells with or without BH-103 treatment.

**Figure 3.**
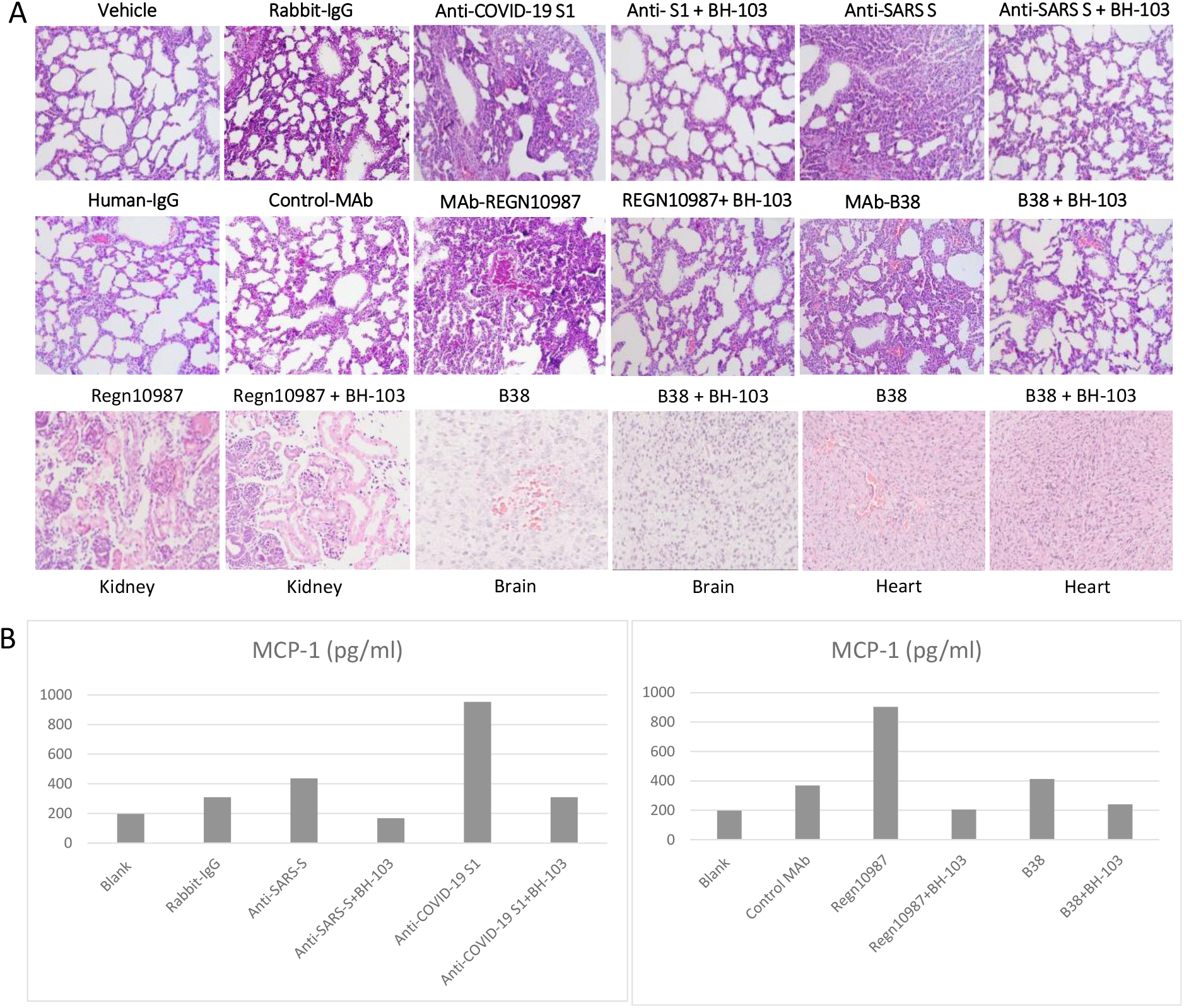
**A**: The representative images of the histological changes of lungs (top 2 rows), kidneys, brains, and hearts (bottom row) from the newborn mouse pups born to the dames injected with pathogenic anti-coronavirus antibodies of anti-Covid-19 S1, anti-SARS S, and human monoclonal anti-COVID-19 S1 antibodies (MAb) REGN-10987 and B38, and control antibodies of human IgG, rabbit IgG and CR3022-b6 (Control MAb); or the dames treated with BH-103 and antibody. **B**: The cytokine levels of MCP-1 in mouse sera from the newborn mouse pups born to the dames with antibody injection alone or the dames treated with BH-103 and antibody.

### Blocking the entry of COVID-19 virus

The angiotensin converting enzyme 2 (ACE2) is an entry receptor for coronaviruses such as SARS-CoV-2. N-acetylneuraminic acid is a component of ACE which is responsible for the attachment of a coronavirus to the receptor on host cells ^9^. We then asked a question: could expression of more sialic acid on cell surfaces increase the receptor amount and infectibility of viruses?

To answer the question, we developed an *in vitro* assay for the viral entry study of the SARS-CoV-2 virus. The human acute promyelocytic leukemia cell line, NB4 cells express ACE2 (The Human Protein Atlas) and was used for the assay. The receptor binding domain (RBD) of the spike protein (S-RBD) of the SARS-CoV-2 virus is responsible for viral entry. The recombinant protein of S-RBD of the SARS-CoV-2 virus (with human Fc as a tag for detection) was purchased (Sino Biological, Beijing) and used for the viral entry assay.

In one test, NB4 cells were cultured with BH-103 as described above, at 50 μg/ml of NANA-Me for overnight, and the sialic acid levels on the cells were determined next day as described above. The results showed that the sialic acid levels on the NB4 cells treated with BH-103 were higher compared with untreated control cells (FIG 2A), indicating the enhancing effect of BH-103 on the sialic acid expression of the NB4 cells.

In a parallel test the NB4 cells, with and without treatment of BH-103, were incubated with the recombinant S-RBD of the COVID-19 virus in ice for one hour and then stained with a fluorescent (PE) labeled anti-human antibody, followed by a flow cytometry analysis. The results showed that the S-RBD bound on the NB4 cells treated with BH-103 was lower compared with that of the control cells without BH-103 treatment (FIG 4B).

**Figure 4.**
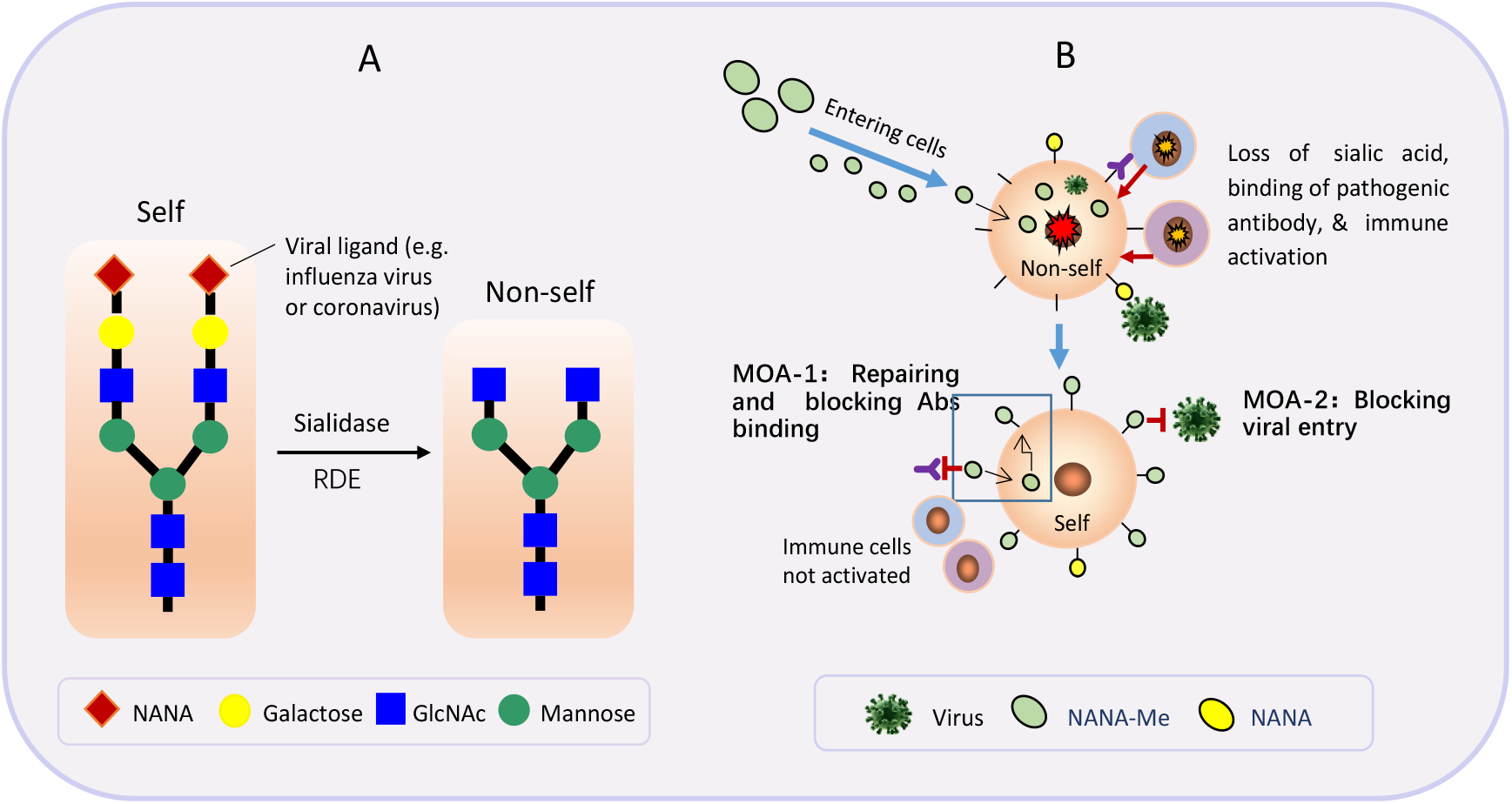
Mechanisms of action (MOA) of N-acetylneuraminic acid methyl ester (NANA-Me). **A**: loss of sialic acid makes self-cells became non-self. **B**. NANA-Me can easily enter cells and participate in the process of sialic acid repairment, and stop ADAA reaction by blocking binding of pathogenic antibody to self-cells (**MOA-1**); and NANA-Me can block viral entry by modifying a viral receptor and dramatically reducing viral binding affinity (**MOA-2)**.

In another test, NB4 cells were treated with neuraminidase, then the sialidase treated NB4 cells were cultured without or with 50 μg/ml of BH-103 for overnight. Next day, the sialic acid levels on the NB4 cells were determined. In a parallel test the binding of the recombinant S-RBD to the sialidase treated NB4 cells with or without treatment of BH-103 were tested as described above.

As shown in FIG 2C, the sialic acid levels of the sialidase treated NB4 cells was lower than that of untreated cells, indicating loss of sialic acid on NB4 cells after being digested with sialidase. The sialic acid levels of the NB4 cells treated with sialidase and BH-103 was higher than that of control cells treated with sialidase alone. Nevertheless, 89% of the binding of the COVID-19 S-RBD to the NB4 cells treated with sialidase and BH-103 was reduced compared to the control cells treated with sialidase alone (FIG 2D). This result indicated that replacement of NANA by NANA-Me induced a structural change or chemical modification of the viral receptor. The data demonstrated that BH-103 not only repaired the missed sialic acid but also significantly decreased the binding affinity of the COVID-19 S-RBD simultaneously.

Taken together, the data indicated that the replacement of NANA by NANA-Me could induce a structural or chemical modification of the viral receptor that significantly decreases the binding affinity of the COVID-19 virus. Therefore, despite the sialic acid expression on the BH-103 treated NB4 cells being higher, the bound S-RBD was significantly lower, especially with the BH-103-treated damaged cells with missing sialic acid. This is because the majority NANA was replaced by NANA-Me on the BH-103-treated damaged cells.

### The prevention and treatment of the adverse reactions of pathogenic antibodies with BH-103

Our recent study has reported a timed-pregnant disease mouse model induced by pathogenic anti-spike antibodies of SARS-CoV-2 and the SARS-CoV viruses^6^.

The mouse model was used in the current study to evaluate the therapeutic effect of BH-103 for the prevention and treatment of the disorders caused by the pathogenic anti-coronavirus antibodies.

Like our previous study, the four pathogenic anti-spike antibodies, the polyclonal rabbit anti-spike (S1) of SARS-CoV-2 (antiCOVID-19 S1), anti-spike glycoprotein (S) of SARS-CoV (anti-SARS S), human monoclonal anti-S1 antibodies of SARS-CoV-2, REGN10987 and B38, were used to create the mouse disease model. The control antibodies included purified IgG of sera from healthy rabbit and human, as well as the monoclonal anti-S1 antibody, CR3022-b6. Two dosages of each purified and endotoxin-free antibody IgG were injected *intraperitoneally (IP)* into timed-pregnant mice of C57BL/6J twice every three days at pregnancy (embryonic) days E15 and E18, respectively. In addition, injection (*IP*) of BH-103 (15 mg/kg) at either day before or the same day of each pathogenic antibody injection were administered.

The frequencies of sickness and death of the fetus and newborn mouse pups are summarized in Table 1. Injection of the four pathogenic antibodies of anti-COVID-19 S1, anti-SARS S, REGN10987 and B38 into pregnant mice induced significant fetal and neonatal death of the mouse pups born to those dames (Table 1). However, the treatment of BH-103 significantly decreased the rates of sickness and death caused by the four pathogenic antibodies (Table 1).

**Table 1.** The sick and death rates of mouse pups born to the dames with the injection of anti-coronavirus antibodies

| Injected IgG of | N = | Sick (%) | Death (%) | Sick+Dead (%) | Odds Ratio | 95% CI | P value |
| --- | --- | --- | --- | --- | --- | --- | --- |
| None (blank) | 7 | 0 | 14.3 | 14.3 | NA | NA | NA |
| Vehicle (saline) | 9 | 11.1 | 0.00 | 11.1 | 0.75 | 0.04-14.6 | 1.00 |
| Healthy rabbit serum | 6 | 0 | 16.7 | 16.7 | 1.2 | 0.06 - 25 | 1.00 |
| Healthy human serum | 10 | 0 | 10.0 | 10 | 0.67 | 0.03 - 13 | 1.00 |
| Anti-COVID-19 S1 | 22 | 28.6 | 28.6 | 57.1 | **9.60** | **1.02 - 90** | **0.04** |
| Anti-COVID-19 S1+BH-103 | 11 | 9.09 | 0.00 | 9.09 | **0.08** | **0.01 - 0.77** | **0.02** |
| Anti-SARS S | 8 | 50.0 | 12.5 | 62.5 | **13.3** | **1.07 - 166** | **0.05** |
| Anti-SARS S+BH-103 | 16 | 6.25 | 6.25 | 12.5 | **0.09** | **0.01 - 0.67** | **0.02** |
| MAb-CR3022-B6 (control) | 9 | 0 | 11.1 | 11.1 | 1.00 | 0.05 - 18.9 | 1.00 |
| MAb-B38 | 24 | 45.8 | 12.5 | 58.3 | **11.2** | **1.20 - 104** | **0.02** |
| B38+BH-103 | 27 | 7.69 | 0 | 3.70 | **0.03** | **0.003 - 0.28** | **< .001** |
| MAb-REGN10987 | 24 | 16.7 | 45.8 | 62.5 | **13.3** | **1.42 - 125** | **0.02** |
| REGN10987+BH-103 | 20 | 5.00 | 0 | 5.00 | **0.03** | **0.004 - 0.28** | **< .001** |

### Histology changes

The tissue sections of lungs, brains, hearts, kidneys, intestines, and livers from the newborn mouse pups were stained with hematoxylin-eosin (HE) for histology evaluation.

Like previous results, acute lung inflammation was observed with the HE stained tissue sections from the mouse pups born to the dames injected with anti-COVID-19 S1, anti-SARS S, REGN10987, and B38 (Figure 2). The lung lesions included pulmonary congestion, alveolar epithelial hyperplasia and thickening, hemorrhage, alveolar atresia, alveolar dilatation, and alveolar fusion (Figure 2A). Infiltration of inflammatory cells at the local lesion areas were also observed. In addition, inflammatory reactions such as acute kidney tubular injury, cerebral and myocardial inflammation and hemorrhage were also observed with the tissues of kidneys, brains, and hearts from the mouse pups born to the dames with the injection of the four pathogenic antibodies (Figure 3). However, the treatment of BH-103 (15 mg/kg), either the day before or at the same time as the antibody injection of the four pathogenic antibodies, significantly decreased the severity of inflammation and injuries of lungs, kidneys, brains, and hearts, compared to the mouse pups delivered to the dames injected with either of the pathogenic antibody alone (FIG 12A). There were insignificant or minor histological changes with the lungs, kidneys, brains, and hearts from the pups born to the dames injected with the control antibodies of CR3022-b6, and the human or rabbit IgGs (Figure 3).

Taken together, the *in vivo* data indicated that BH-103 is effective for the prevention and treatment of the inflammation and injuries of multiple organs caused by pathogenic antibodies of the COVID-19 and SARS-CoV viruses.

### BH-103 significantly reduced cytokine production

The sera from newborn pups as mentioned above were tested for inflammatory cytokines of MCP-1, TNF-α, IL-4, IL-6 and IL-10 using a 5-plex multiplex Luminex assay kit (Millipore) according to manufacturer’s instruction. The results are summarized in FIG 3B. The antibodies of anti-COVID-19 S1 and REGN10987 induced significantly higher levels of MCP-1 (Figure 3B). Nevertheless, the treatment of BH-103 significantly reduced the cytokine levels of MCP-1 (P < 0.001), compared to the treatment with the pathogenic antibody alone. The results were consistent with the results of the sickness and death rates (Table 1) and the histology changes (Figure 2). The data further indicated that BH-103 is effective in preventing or treating cytokine storms caused by pathogenic antibodies. The levels of other cytokines were not significantly elevated, probably due to the undeveloped immunity of the newborn mouse pups.

## Discussion

### The novel mechanism of action (MOA)

#### MOA-1: Repairing and blocking antibody binding

Beside their major function as biological masks, sialic acids also act as viral receptors^8^. Certain viruses, including coronaviruses, contain receptor-destroying enzymes (RDE) that cleave a terminal sialic acid from the cellular surface^9^. During an infection of a virus with RDE, the terminal sialic acids on cell surfaces can be removed, and the cellular structures become “non-self”^8^ (Figure 4A). Our recent study has shown that damaged lung epithelium cells with missing sialic acid are particularly vulnerable to pathogenic antibodies of the COVID-19 virus^6^. Thus, repairment of the missing sialic acid can convert the sick cells (non-self) back to normal (self) and stop the subsequent autoimmune reactions (ADAA) caused by pathogenic antibodies. These autoimmune reactions are not minor side effects of the infection, but rather can be the main contributors to the severity of an infection and can lead to serious clinical conditions including ARDS, cytokine storms, and even death.

So far, there have been no reports that uses analogs or derivatives of sialic acid as repairers or modifiers of viral receptors on host cells, probably because most anti-viral products have focused on targeting viruses. Here, we explore a new application of an analog of N-acetylneuraminic acid (NANA), N-acetylneuraminic acid methyl ester (NANA-Me), for the prevention and treatment of serious symptoms of COVID-19 infection, through a unique MOA that is being reported for the first time.

The structure of NANA-Me is close to the structure of NANA. Unlike NANA, NANA-Me can easily penetrate the cell membrane, enter cells, and participate in the sugar chain synthesis process. Therefore, NANA-Me is a good candidate for the repairment of missing sialic acid on the damaged lung epithelium cells or other cells such as inflammatory cells (Figure 4B, MOA-1). The structure recovery of the damaged cells can reduce the binding of pathogenic antibodies and stop the ADAA reactions of immune system against the damaged or inflammatory cells. These MOA of NANA-Me is supported by the *in vitro* and *in vivo* data of the current study. The *in vitro* data showed that NANA-Me can enhance or repair NANA expression on lung epithelium A549 cells (Figure 1). The *in vivo* results showed that BH-103 effectively prevented or treated the adverse reactions of pathogenic anti-spike antibodies of COVID-19 and SARS-CoV viruses (Table 1). Because the anti-spike pathogenic antibodies are also inducible by COVID-19 vaccines, BH-103 can be effective for preventing and treating the adverse reactions of COVID-19 vaccines.

#### MOA-2: Receptor modification

Many microbes bind to mammalian tissues by recognizing specific saccharide ligands. For example, sialic acid moieties on glycoproteins are critical receptor determinants, and are related to viral attachment of influenza viruses or coronaviruses (Stevens et al., 2006, Huang et al., 2015). Therefore, saccharides and saccharide mimetics can be used to block the initial attachment of microbes to cell surfaces or block their release, thus preventing and suppressing infection (anti-adhesive). Examples of such applications include milk oligosaccharides that are believed to be natural antagonists of gastrointestinal infection in infants, and polymers that will block the binding of viruses ^11^.

In addition, chemical modification of sialic acids can strongly influence their properties, in particular ligand functions^11^. The hydroxyl groups present in both monosaccharides and oligosaccharides can be chemically modified without affecting the glycosidic linkages ^11^. Natural products containing partially methylated glycans are known and a number of methyltransferases have been identified ^11^. For example, O-methylation can hinder or even prevent hydrolysis of the glycosidic bond by sialidase^11^. Nevertheless, most derivatives of sialic acid, including some methyl esters of NANA, are widely used for competitive inhibitors of neuraminidase of bacteria and viruses (e.g. influenza viruses)^12^. The most well-known such example is *Tamiflu*.

NANA-Me can act as a donor of methyl organ. The replacement of NANA by NANA-Me could cause a structural or chemical modification of the viral receptor that can significantly decreased the binding affinity of the related viruses including the SARS-CoV-2 (MOA-2, Figure 4B) (Figure 2).

### Why in combination with NANA?

One weakness of NANA-Me is that it is unstable and easily hydrolyzed. This weakness makes it hard for NANA-Me to be a good therapeutic product. Our study found that the stability of NANA-Me was improved when it was in a composition comprising NANA and under a condition of pH 4.0-5.5 (unpublished data, patent pending). Therefore, we designed the formulation of BH-103 consisted of NANA-Me and NANA with a ratio of NANA-Me:NANA at 2:1 (pH 4.43). Our *in vitro* and *in vivo* data showed that BH-103 has overcome this shortcoming and significantly improved the efficacy of NANA-Me. Thus BH-103 can be a good therapeutic medicine for the disorders caused by pathogenic antibodies of viral infections such as COVID-19.

### Advantages of drugs targeting viral receptors

The results of this study indicate that NANA-Me (or BH-103) can chemically modify the sialic acid of the SARS-CoV-2 viral receptor and block viral entry into host cells (MOA-2, Figure 2 and Figure 4B). Based on the MOA-2, N-acetylneuraminic acid methyl ester or BH-103 has the potential to prevent the COVID-19 infection by blocking viral entry into host cells, and treat the infection by blocking viral spread into new cells. Future prospective *in vivo* studies are needed to confirm this function of NANA-Me.

Given the similarities in receptor usage, sialic acid is a receptor component for not only coronaviruses, but also other viruses such as influenza viruses. The receptor modification and entry blockage by NANA-Me should be widely effective for the prevention and treatment of other infections caused by other viruses using sialic acids as receptors, particular a highly pathogenic influenza infection. As supporting evidence, our previous studies showed that formulations consisting of NANA-Me and NANA were effective for prevention and treatment of influenza infections ^13^.

There are a couple of advantages with therapeutics targeting viral receptors: 1) broad spectrum of efficacy. Unlike the multiple and complicated viral strains, viral receptors on host cells are limited. Thus, a therapeutic capable of modifying receptors can be effective to all the viruses and viral strains sharing the receptor. 2) anti-mutation. Effective modification of a viral receptor can block the attachment of all viruses using that receptor, regardless their strains (including mutated variants).

### Highlights of BH-103

Compared to other anti-viral drugs which target virus, BH-103 has the advantages below: Firstly, BH-103 acts through a unique MOA of interrupting ADAA caused by pathogenic antibodies from all viruses capable of inducing such antibodies, for example highly pathogenic coronaviruses such as COVID-19 virus and influenza viruses.

Secondly, BH-103 is broadly effective for the ADAA-based diseases caused by pathogenic antibodies. Those diseases include infectious diseases, complications and sequela of infections including COVID-19 long haulers, cytokine storm and cytokine release syndrome (CRS), adverse reactions of vaccines, infection-relating autoimmune diseases, and other possible disorders inducible by pathogenic antibodies. The ADAA-based diseases further include abortion, postpartum labor, still birth and neonatal death of pregnant females, caused by an infection or by a vaccine.

Thirdly, BH-103 has an excellent safety profile. A 7-day multiple dosing toxicity study with rats and dogs showed that no signs of toxicity was observed with BH-103 at the highest dose of 750 mg/kg (unpublished data).

Fourthly, no drug resistance. BH-103 is resistant to viral mutations since it targets a host receptor instead of a virus.

Fifthly, BH-103 is easy to produce and large amounts of it can be produced in a short amount of time. This will be helpful in fulfilling the large increase in demand for effective therapeutics experienced during a pandemic, such as the current COVID-19 pandemic.

**In summary**, our recent study revealed “pathogenic antibodies” in viral infections. These pathogenic antibodies act through a novel mechanism of action that we have termed “Antibody Dependent Auto-Attack” (ADAA). The current study reports a formulation (BH-103) that is designed to interrupt the ADAA reactions through blocking pathogenic antibody binding. Therefore, BH-103 can be broadly effective for the diseases caused by pathogenic antibodies, such as serious COVID-19 conditions including ARDS, cytokine storms, and death. Further, BH-103 can be effective for infection-related autoimmune diseases, including those experienced by COVID-19 long haulers. Importantly, BH-103 can prevent and treat the adverse reactions of vaccines including COVID-19 vaccines. This will be particularly helpful to promote the global COVID-19 vaccination and to control the pandemic as soon as possible. Therefore, BH-103 will provide an effective resolution for the urgent clinic unmet of the COVID-19 pandemic and the other possible pandemic emerging in future.

## Methods

### Compounds

Compounds of N-acetylneuraminic acid (NANA) and N-acetylneuraminic acid methyl ester (NANA-Me) were purchased (Jun Kang Biotech, Guangzhou). NANA-Me comprises the formula of C12H21NO9, and the molecular structure of:

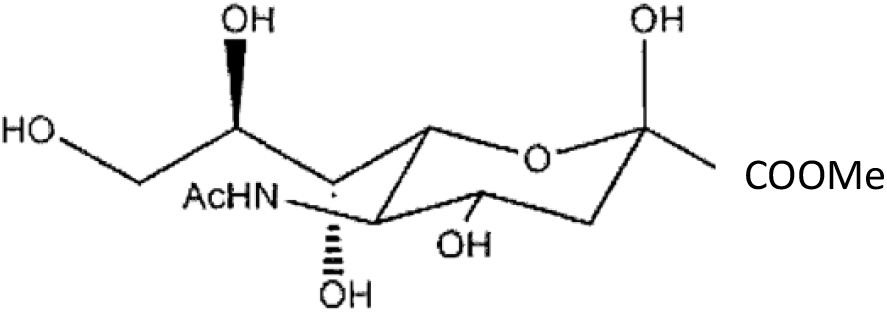

The compound purity was 97% for NANA, and 98% for NANA-Me. A formulation consisted of 0.5 gram of NANA-Me, 0.25 gram of NANA, and 0.175 gram of sodium citrate was made and named BH-103. A solution of BH-103 at 10 mg/ml (pH 4.43) was made, aliquoted and stored at -80°C. Freshly thawed BH-103 solution (pH 4.43) was used for each experiment.

### Production of human anti-SARC-CoV-2 spike monoclonal antibodies

Previously reported naturally occurring human monoclonal antibodies of B38 ^14^, REGN10987 ^15^, and CR3022-B6 ^16^, specific to the receptor binding domain (RBD) of the spike protein one (S1) of the SARS-CoV-2 virus were re-produced as previously reported (HuaAn McAb Biotechnology). Removal of endotoxin from antibody solutions was performed as previously reported.

### Other antibodies

The rabbit polyclonal antibodies specific for the recombinant spike one (S1) proteins of the SARS-CoV-2 virus, the recombinant spike proteins of SARS-CoV virus, were purchased from Bioss Antibodies (Beijing).

### S-RBD of SARS-CoV-2

The recombinant protein of the receptor binding domain (RBD) of the spike protein (S-RBD) of the SARS-CoV-2 virus (with human Fc as a tag for detection) was purchased from *Sino Biological* (Beijing).

### Digestion of A549 and NB4 cells with neuraminidase

The cells of human lung epithelium cell line A549 (ATCC), and human acute promyelocytic leukemia cell line NB4 (ATCC) were washed once with the digestion buffer for neuraminidase (Roche), the cells were centrifuged at 2000 rpm for 5 minutes, and the supernatant was discarded. The digestion buffer was consisted of 50 mM of sodium acetate, 3% BSA, and 2 mM of CaCl2 in PBS (pH5.5-6.0). The cells were resuspended with the digestion buffer and divided to multiple tubes, each tube contained about 10^6^ of cells in 200 μl of the digestion buffer containing 50 μU of neuraminidase (Roche). The tubes were incubated at 37°C for one hour. Then the cells were washed with PBS once and proceeded for flow cytometry assay. A549 or NB4 cells without the treatment of neuraminidase were used as controls.

### Binding of S-RBD to NB4 cells and flow cytometry

NB4 cells with or without the treatment of neuraminidase or BH-103 were tested for the binding of the S-RBD of the SARS-CoV-2 virus as mentioned above. Each 2×10^5^ of cells in 200 μl of 1%BSA-PBS were incubated in ice with the S-RBD at a concentration of 5 μg/ml for one hour, washed once with PBS. The cells were resuspended with 200 μl of 1%BSA-PBS and 0.25 mg/ml of PE-labeled goat anti-human IgG (Abcam), incubated in ice for 30 minutes, and washed once with PBS. The cells were resuspended with 200 μl of PBS and detected with a flow cytometer of Accuri 6 (BD Biosciences). The fluorescent labeled wheat germ agglutinin (WGA), which specifically binds to sialic acid (Vector), was used as a secondary reagent for determine the sialic acid level of cells as needed.

### A timed-pregnant mouse model

A timed-pregnant mouse model without viral infection was developed using the anti-coronavirus antibodies as previously reported. The animal experiments were performed at the Center of Laboratory Animals of Hangzhou Normal University. The protocol for the animal experiment was approved by the Laboratory Animal Welfare and Ethics Committee of Hangzhou Normal University. The CALAS No. is 2020244. The animal experiments were performed three times.

### Animal

SPF-grade C57BL/6J pregnant mice at pregnancy (embryonic) day E13-E14 were purchased from Shanghai SLAC Laboratory Animal Co., Ltd. The animals were randomly divided into groups as needed, two pregnant mice for each group at every experiment.

### Antibody injection

The purified and endotoxin-free IgG of the anti-coronavirus antibodies used for the animal model include rabbit polyclonal anti-COVID-19 S1, anti-SARS-CoV S, and the human monoclonal anti-COVID-19 S1, B38^14^, REGN10987^15^, and CR3022-B6 ^16^. Two dosages of each antibody IgG were injected *intraperitoneally (IP)* into timed-pregnant mice once every three days at pregnancy (embryonic) day of E15 (about 26-28 g) and E18 (about 30-32 g) respectively. For each polyclonal antibody, 50 μg (microgram) for the first dose (about 2.0 mg/kg) and 60 μg for the second dose (about 2.0 mg/kg) were administrated. For each monoclonal antibody, 40 μg for the first dose (about 1.5 mg/kg) and 50 μg for the second dose (about 1.5 mg/kg) were administrated. In addition, two dosages of BH-103 injection (*IP*) at either day before or the same day of each antibody injection of the 4 pathogenic antibodies (anti-COVID-19 S1, anti-SARS-CoV S, REGN10987, and B38) were administrated, respectively.

The body weight of the pregnant mice was measured every day before and after the antibody injection. The mouse pups were born at about E20-E21 and the healthy status including clinical signs of the newborn mouse pups were observed and recorded. The experiment was ended at day 1-2 post birth.

### Sample collection

At the end day of the experiment, the blood samples were collected from newborn mouse pups, incubated at 4°C for overnight, centrifuged at 3000 rpm for 5 minutes, and the supernatant was transferred to a new tube. The isolated sera were stored at -80°C for cytokine detection. Lungs, hearts, brains, kidneys, livers, and intestines were collected from at least 3 mouse pups, fixed in formalin for 48-72 hours, went through gradient alcohol dehydration and embedded in paraffin, and tissue sections were processed.

### HE and immunofluorescent staining

The mouse tissue sections of lungs, hearts, brains, kidneys, livers, and intestines were dewaxed and stained with hematoxylin-eosin (HE) for histology evaluation.

## Acknowledgement

The study was supported by funding from Hangzhou HuaAn PuChu Investment Limited Partnership.

## Author Contributions

H.W. conducted the overall study direction including experiment design, data analysis, and manuscript writing; Q.C. and D.L. conducted the *in vitro* cellular analysis, IgG isolation and removing endotoxin; X.W. conducted the animal experiments, sample collection and preparation of tissue section; Y.Z. conducted the process of tissue sections and HE staining, Y.C. conducted funding acquisition, resources, and supervision; X.L. conducted antibody production, data analysis and discussion.

## Competing Interests

H.W. is the shareholder of Huirui Biopharma, X.L. and Y.C. are shareholders of HuaAn McAb Biotechnology. There are patent applications pending related to this work.

## Notes

https://www.ncbi.nlm.nih.gov/Structure/pdb/7BZ5/6XDG/7KZB

### Summary of Updates

The Table 1 was replaced.

